# Mechanics of snake biting: Experiments and modelling

**DOI:** 10.1101/815407

**Authors:** Lakshminath Kundanati, Roberto Guarino, Michele Menegon, Nicola M. Pugno

**Author notes:** École Polytechnique Fédérale de Lausanne (EPFL), Swiss Plasma Center (SPC), CH-5232 Villigen PSI, Switzerland. PAMS foundation, Tanzania.

## Abstract

Among all the vertebrates, snakes possess the most sophisticated venom delivering system using their fangs. Fangs of many animals are well adapted to the mechanical loads experienced during the functions such as breaking the diet and puncturing the skin of the prey. Thus, investigation and modelling of puncturing mechanics of snakes is of importance to understand the form-function relationship of the fangs and tissue-fang interactions in detail. We have thus chosen fangs of two snake species i.e. viper (*Bitis arietans*) and burrowing snake (*Atractaspis aterrima*), with different shape and size and performed insertion experiments using tissue phantoms. Our results showed that both the species have similar mechanical properties but there was a difference in the insertion forces owing to the difference in shape of the fang. Also, our modelling of the fang-tissue interactions predicted some parameters close to the experimental values. Thus, our study can help in the development of bioinspired needles that can potentially have reduced insertion forces and less damage to the tissue.

## 1. Introduction

Biological structures like teeth are known to adapt well to mechanical loading conditions (Meyers et al., 2013). Among all the vertebrates, snakes possess the most sophisticated venom delivering system using their fangs (Jackson, 2002). Fangs are the special teeth that are used to inject venom in to prey by many species, using a venom canal that runs through (Zahradnicek et al., 2008). Their long tubular fangs facilitate injection of venom deep into the skin of the prey (Jackson, 2003). In order to cut through the prey tissue, the material of the fang must be of similar or of stiffer material (Politi et al., 2012). As mentioned, teeth and fangs of many animals are well adapted to the mechanical loads experienced during the function of either breaking the diet or puncturing the skin of the prey. Spider fang was observed to have a design with fine mechanical tuning of properties at different locations for easy piercing, reducing wear and withstanding stresses (Politi et al., 2012). Changes in the shape and size of the fang during evolution could have occurred also with the goal of injection system (Kardong, 2016). Investigating mechanics of puncturing by various snakes is of importance to understand the form-function relationship of the fangs.

Understanding the tissue-fang interaction is important to get an overall idea of the biomechanics of insertion during biting. There are many earlier studies which addressed the needle-tissue interactions with the goal of designing needles that induce minimum pain (O’Leary et al., 2003; Shergold and Fleck, 2004). Modelling of the forces involved in puncturing is done by separating the contributions coming from the stiffness of the tissue or phantom material, the cutting force or piercing force during the initial phase of insertion and finally the frictional force between the needle surface and the substrate (Okamura et al., 2004). Most of these studies are based on standard suture needles or needles specially developed for the percutaneous use. There are a very few studies which directly used the fang or piercing organ of an animal to study the interaction (Matushkina and Gorb, 2007; Zhao et al., 2015). Using natural piercing organs also present more challenges but the experiments helps in understanding the interaction in a better way.

The goal of our study is threefold. Firstly, to perform piercing experiments on a substrate that has material properties like Young’s modulus close to that of human or animal skin, to understand the mechanics of fang insertion. Secondly, to determine the mechanical properties of the fangs for comparison. Finally, to model analytically the insertion process at various stages and to validate the model with the experimental data. We used fangs of two snake species i.e. viper (*Bitis arietans*) and burrowing snake (*Atractaspis aterrima*), with different shape and size. This viper species preys on mammals, birds, reptiles and humans, and is known for causing death to both human and animals at times. They have long fangs which can rotate and are hollow (Cundall, 2015). On the other hand, burrowing snake has relatively shorter fangs and preys upon relatively small animals as compared to the viper species. In order to understand the role of speed on the insertion force, we have performed experiments at three piercing speeds. This study would help in understanding the mechanics of fang insertion during biting and can be extended to aid in the development of needle design in biomedical applications.

## 2. Materials and Methods

### 2.1. Microscopy

Images of the fangs are taken using an optical microscope (Lynx LM-1322, OLYMPUS) and a CCD camera (Nikon) attached to the microscope. The dimensions from the optical images are quantified using the standard calibration.

Scanning Electron Microscopy (SEM) was performed on the prepared fangs post mechanical tests. They are carefully mounted on double-sided carbon tape, stuck on an aluminium stub followed by sputter coating (Manual sputter coater, Agar scientific) with gold. A SEM (EVO 40 XVP, ZEISS, Germany) was used with accelerating voltages between 5 and 20 kV. ImageJ software was used for all the dimensional quantification reported in this study (Abràmoff and Magalhães, 2004).

### 2.2. Gel preparation

Food grade gelatine was used to make the phantom gels. The plate-like gelatine was broken into pieces and measured in the weighing balance to mix right proportions (1 g gelatine in 5 ml of water). The pieces of gelatine were soaked in water for 10 minutes and later thoroughly mixed in water with a temperature around 80°C. The gel was then poured in Teflon moulds for curing and placed inside a refrigerator for gelation. The cured gels were carefully taken out of the moulds and used for experiments.

### 2.3. Compression testing

We have estimated the bulk mechanical properties of the gelatine hydrogels using rectangular blocks (15 mm × 10 mm × 9.5 mm). We applied force on the gels (number of samples = 4) using a flat platen at a rate of 0.01 mm/sec, using Messphysik MIDI 10 (MESSPHYSIK, Germany) Universal Testing Machine and the forces were recorded using transducer of (LEANE Corp., ±2 N). The initial linear region of the stress-strain curve is used to estimate the Young’s modulus of the material.

### 2.4. Wire cutting

We used wires of three different diameters (0.18, 0.4 and 0.8 mm) to determine the fracture toughness of the gels (2 samples for each diameter) which are used in the piercing experiments. The wires were kept taught between two points on the custom made set up and are pushed in the gel along the width of the gel block. The gels used in these experiments were prepared as mentioned in the corresponding section. The wires were pushed at a rate of 0.1 mm/s through the gels.

### 2.5. Piercing force experiments

The fangs were fixed in resin at the base of the fang to aid in holding the samples without causing damage. Piercing experiments were performed using a Messphysik MIDI 10 (MESSPHYSIK, Germany) Universal Testing Machine and the forces are obtained using transducer of (LEANE Corp., ±0.25N). Specimens (3 piercing at each rate) were pierced in displacement-control mode at different rates (0.01, 0.1 and 1 mm/s) until the straight portion of the fang was inserted into the gel. The various stages of gel deformation and needle insertion and retraction are depicted in the schematic (Figure 1).

**Figure 1.**
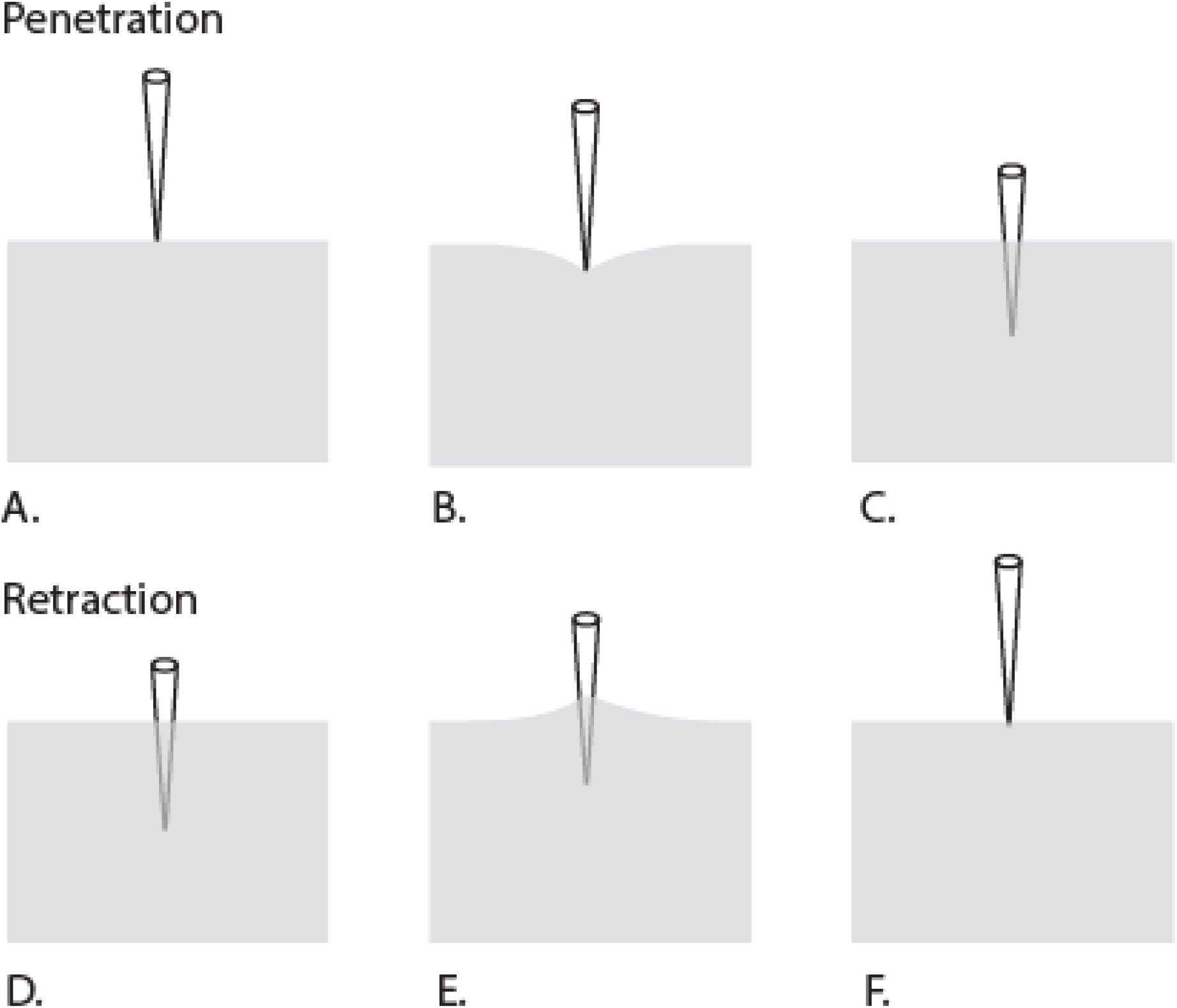
Different stages of needle movement during penetration and retraction.

### 2.6. Nanoindentation

All fangs are embedded in a resin and polished using a series of 400, 800, 1200, 2000 and 4000 grade sand papers. Finally, the sample was polished using a diamond paste of particle sizes in the range of 6 μm and 1 μm, to obtain a surface of minimal roughness. The material properties of fangs are then determined using Nanoindentation technique. We used Berkovich indenter to perform nanoindentation experiments with a maximum load of 30 mN on the polished cross-sectional surface of the tooth samples. We used a matrix format (3×3) to perform a total of 18 indentations with 9 at each location with a prescribed distance between them. A Poisson’s ratio of 0.31 was used for estimating the Young’s modulus.

## 3. Results and Discussion

### 3.1. Morphology and properties of fangs

The shape and tip morphology are determined to see their effects on the insertion force during the experiments of the two selection snake species, viper (*Bitis arietans*) and burrowing snake (*Atractaspis aterrima*) (Figure 2A & 2D). The optical images show that the fang of the viper is longer and has more curvature as compared to the burrower (Figure 2B & 2E). Scanning electron images of the fang tips show that the tip sharpness is almost similar in both the species (Figure 2C & 2F).

**Figure 2.**
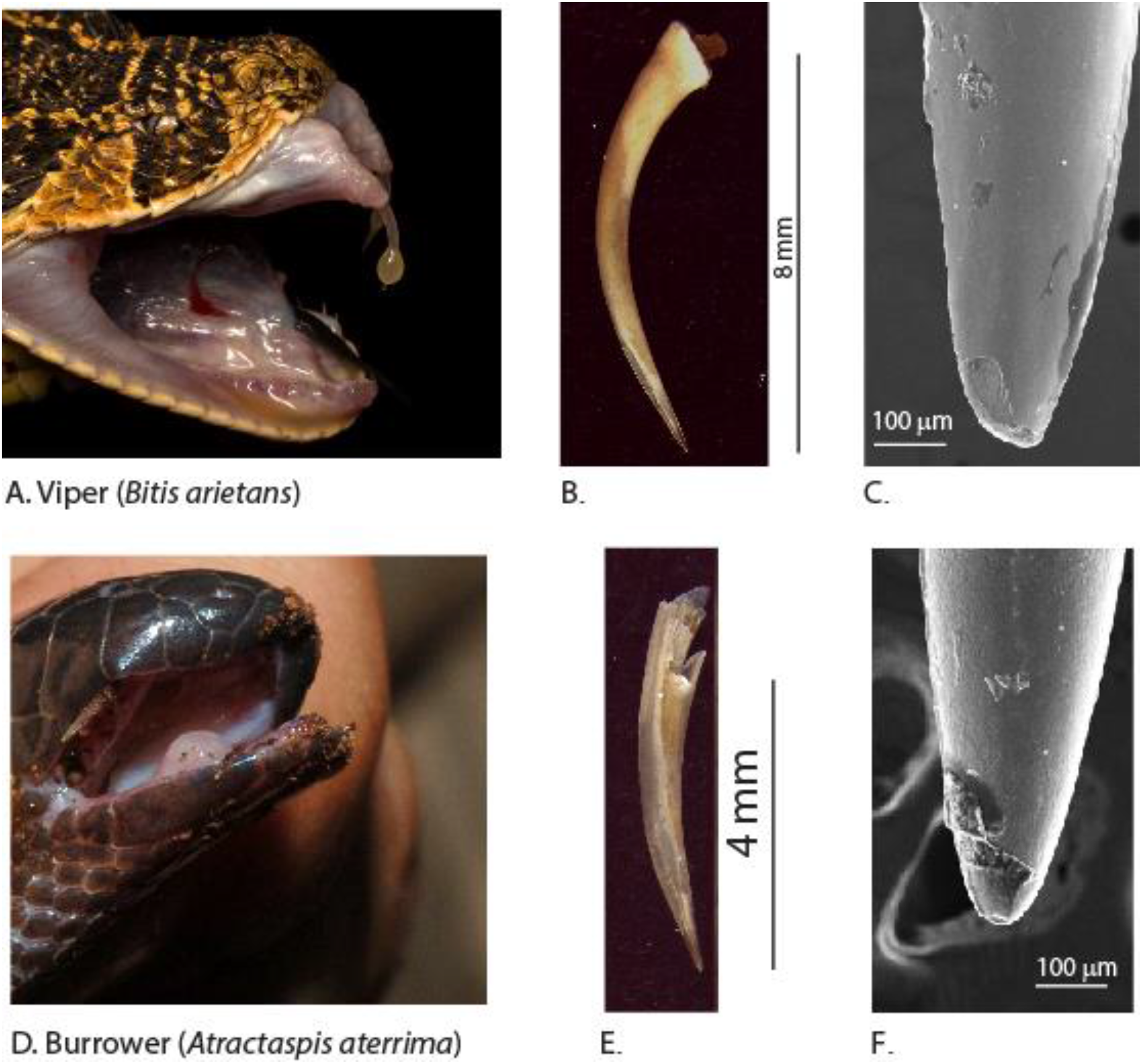
**Images of viper snake A.** head and mouth parts (Image: ©Tyrone Ping (www.tyroneping.co.za)). **B.** Fang **C**. SEM image of it. **Images of burrower snake D**. head and mouth parts **E**. Fang **F.** SEM image of it.

The mechanical properties of the fangs were determined by performing nanoindentation at different locations on the polished sample surfaces (Figure 3). The elastic modulus and hardness of the fangs of the burrower and viper are similar (Table 1). There are also no significant differences between the tip region and the base region of the fang. These values are in agreement with the Young’s modulus (15.3-24.6 GPa) of a few snake species (Jansen van Vuuren et al., 2016)

**Figure 3.**
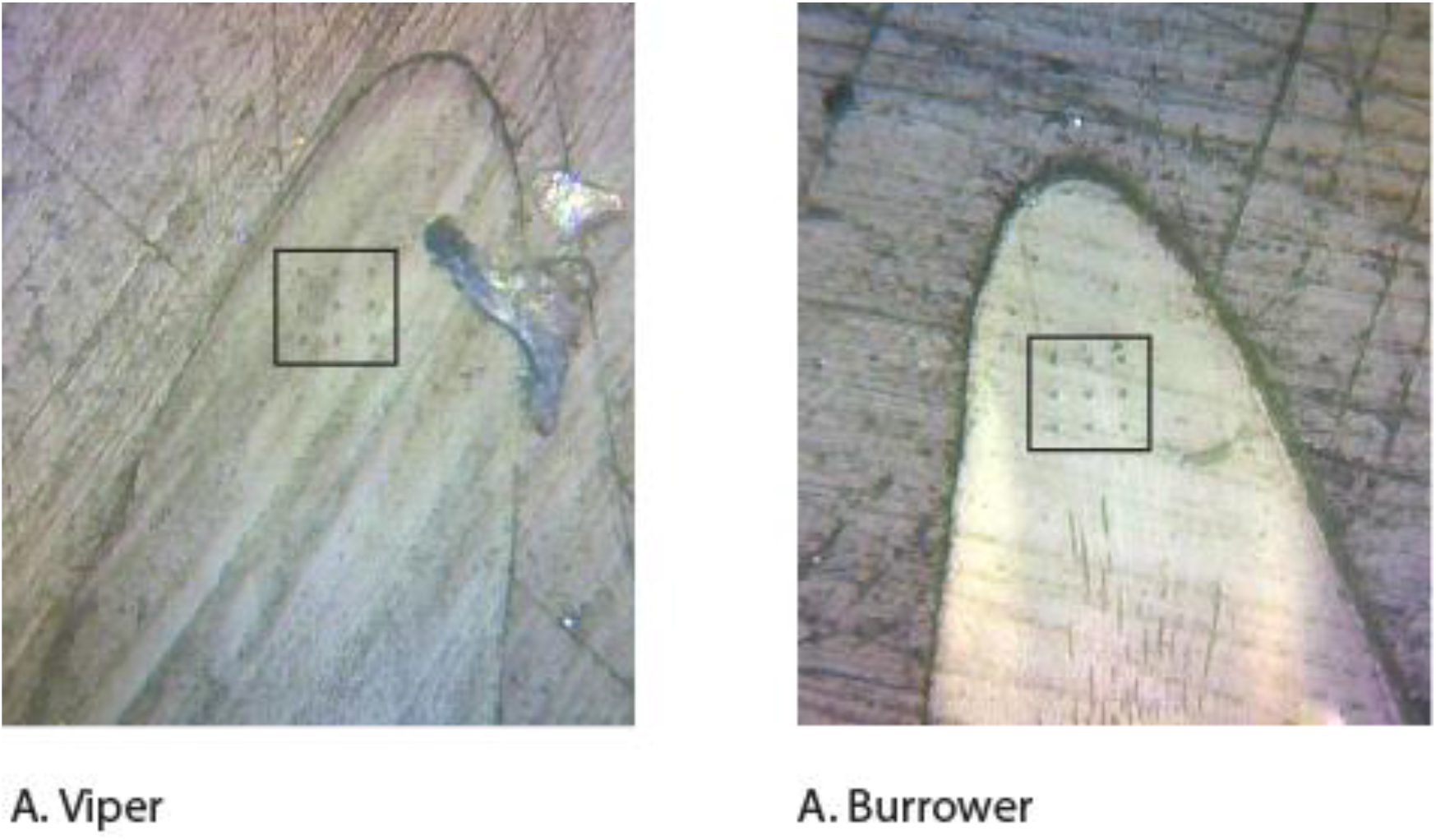
The indentation marks are clearly visible on the polished cross-section of the samples.

**Table 1.** Elastic modulus and hardness of the fangs

| Sample | Region | Elastic modulus (GPa) | Hardness (GPa) |
| --- | --- | --- | --- |
| Viper | Tip | 17.8±2.3 | 0.71±0.06 |
|  | Base | 19.2±0.4 | 0.75±0.01 |
| Burrower | Tip | 18.9±4.7 | 0.92±0.05 |
|  | Base | 18.7±2.0 | 0.86±0.03 |

#### 3.1.1. Compression testing and piercing testing

The stress-strain curves from the compression testing of the gelatine hydrogels showed good repeatability (Fig. 4). The Young’s modulus of the gelatine hydrogels was measured to be 380±65 kPa. Using gels of same composition, the force-displacement curves are obtained from the piercing tests performed at different rates. The curves resembled standard piercing tests with a linear increase in force with increase in displacement followed by a sudden drop in force and then a gradual increase (Fig. 5). Using the curves, the force and depth values just before piercing are obtained and compared from different rates. We also measured the force value drop after piercing. The piercing forces of burrowing snake fang are 23±9, 23±3 and 27±1 mN for rates of insertion 0.01, 0.1 and 1 mm/s, respectively. While the values of substrate surface deflection at the time piercing varied a bit. The piercing forces of viper snake fang are found to be 36±6, 37±8 and 37±8 mN for rates of insertion 0.01, 0.1 and 1mm/sec respectively. In both the species, the piercing force and substrate surface deflection at the time piercing did not vary significantly with increase in speed of insertion. We also observed that the fangs experience a negative pull while the fangs were being retracted because of frictional and adhesive effects at the fang-gel interface (Fig. 5). The retraction forces were observed to be lower when the speed of insertion was lower (Table 2).

**Figure 4.**
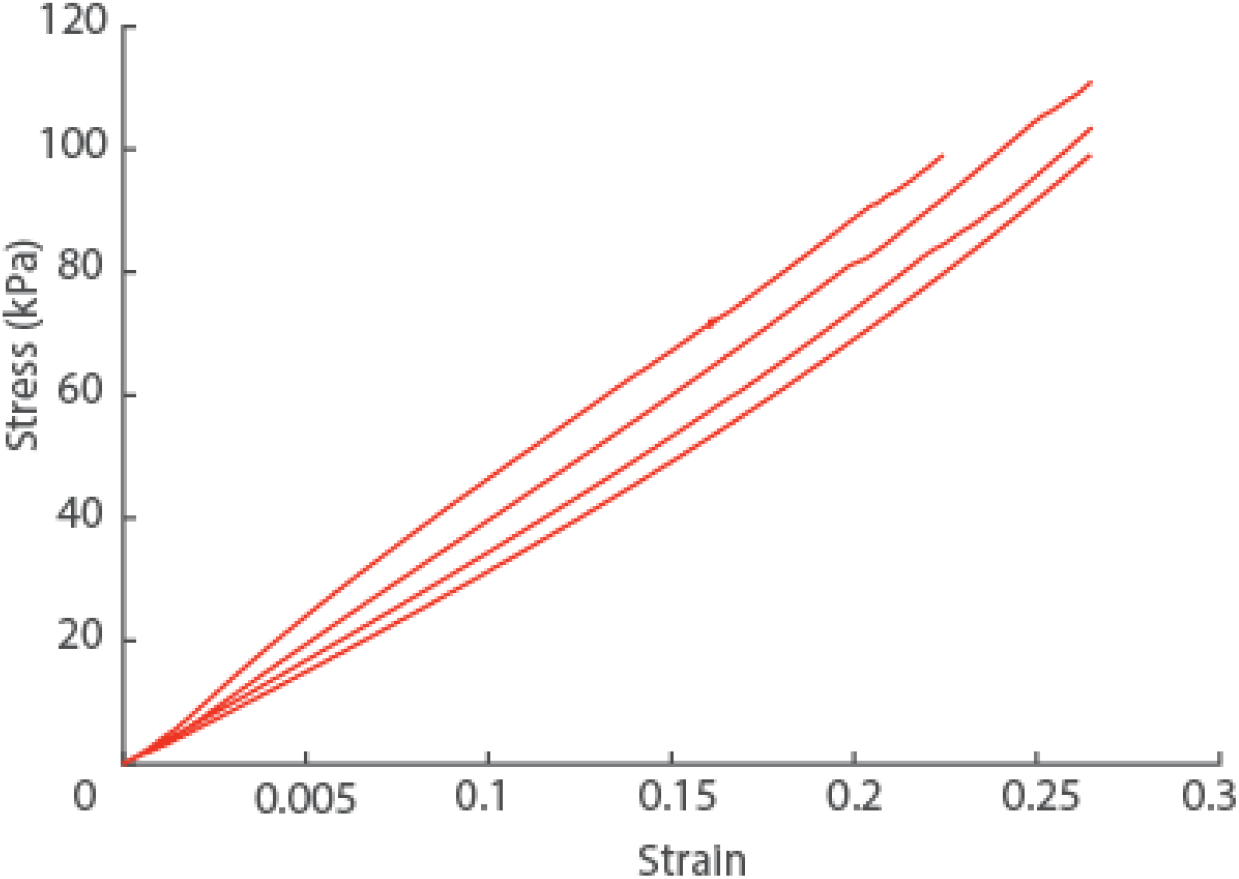
Stress-strain curves from the compression testing of the gelatine blocks.

**Figure 5.**
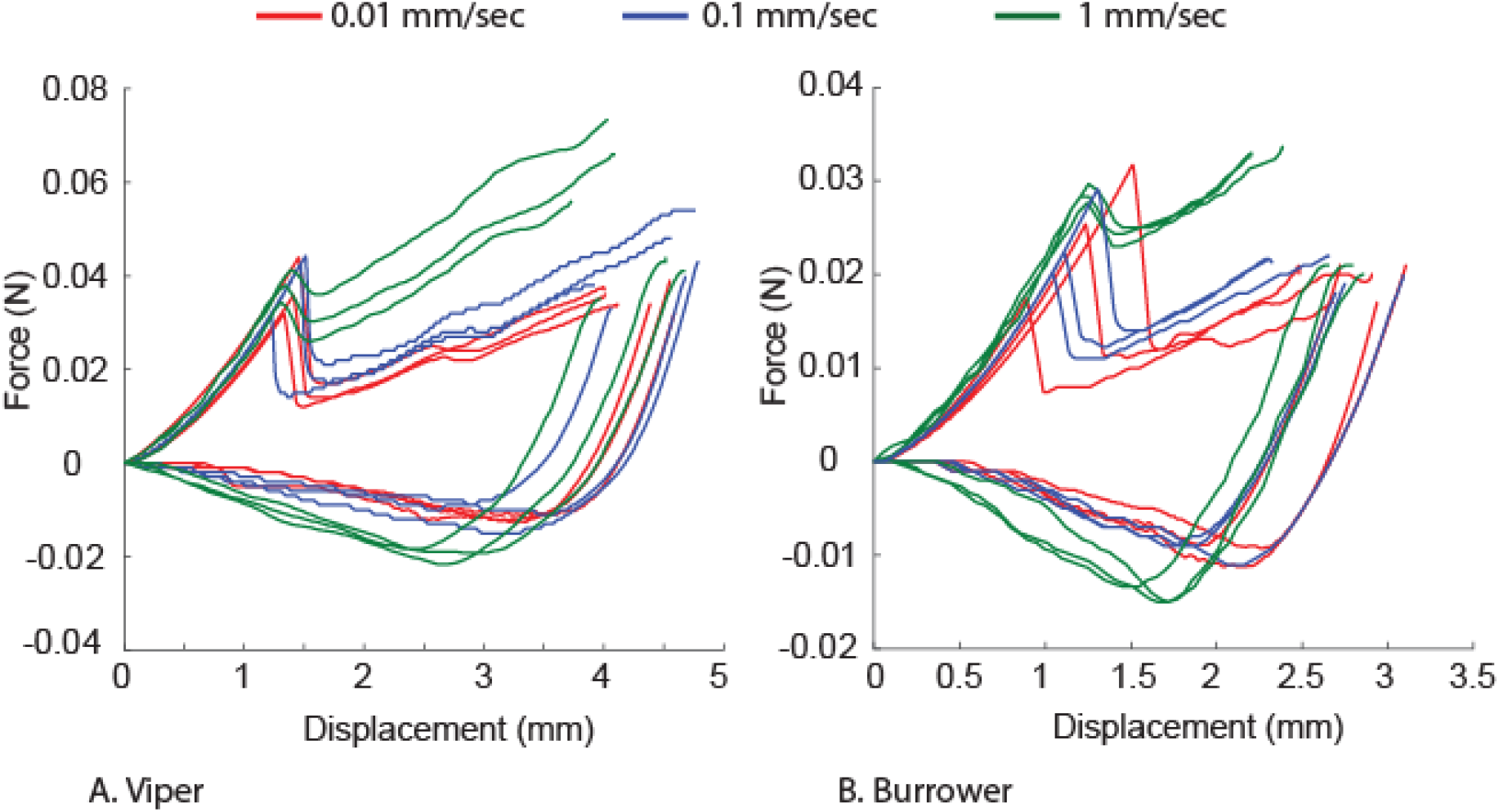
Force-displacement curves during insertion and retraction at different rates. A) Viper B) Burrower snake.

**Table 2.**
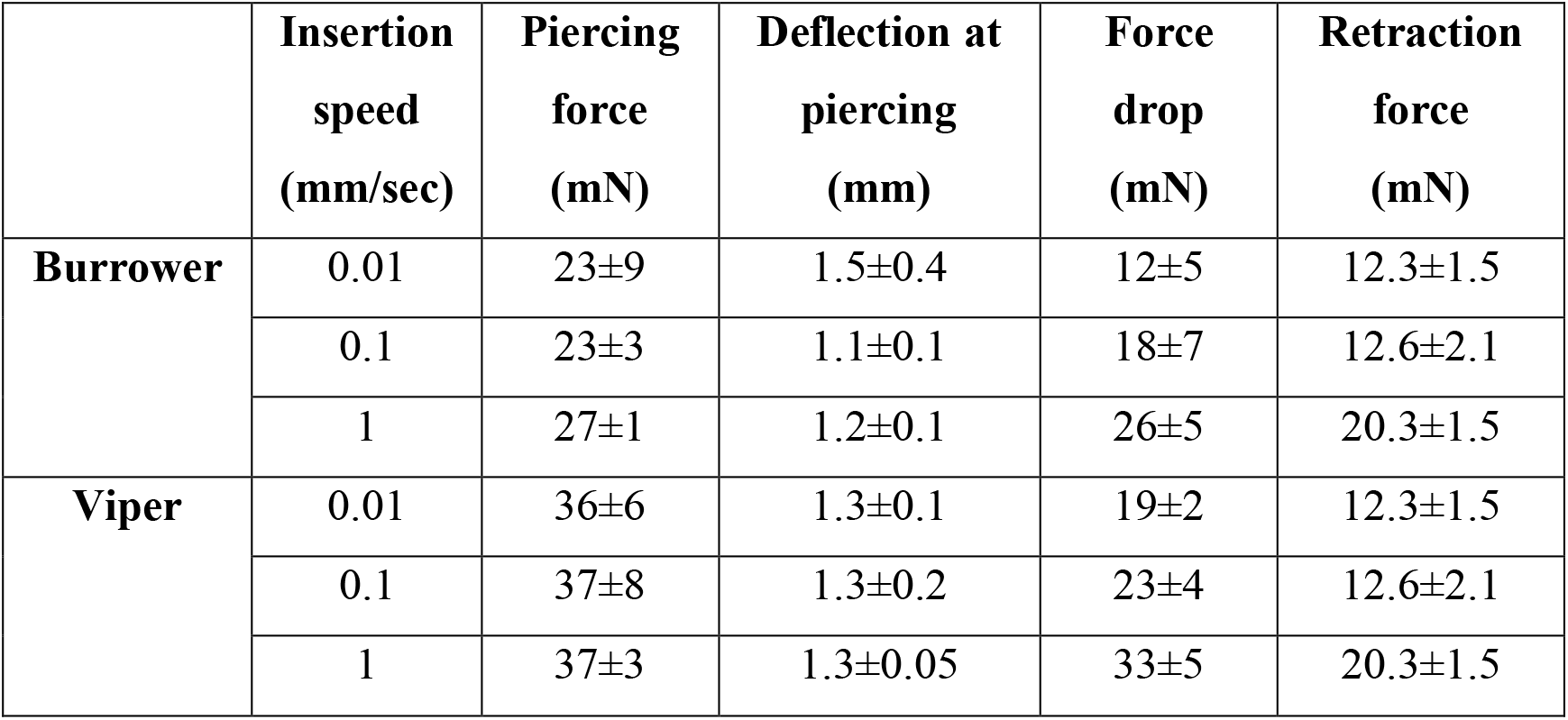
Piercing force of burrowing snake and Viper snake

#### 3.1.2. Wire cutting tests

Wire-cutting tests were performed to determine the Griffith critical energy release rate (*G_c_*). The average cutting force values were determined from the force-insertion curves. These values were divided by the corresponding breadth of gel block and are plotted against the diameter of the corresponding wire diameters. A straight line is fit to the data points using the equation below (Kamyab et al., 1998) and the intercept of the line-fit represents the Griffith critical energy release rate (Fig. 6).

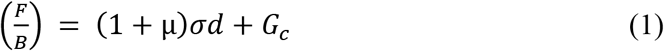

where,

**Figure 6.**
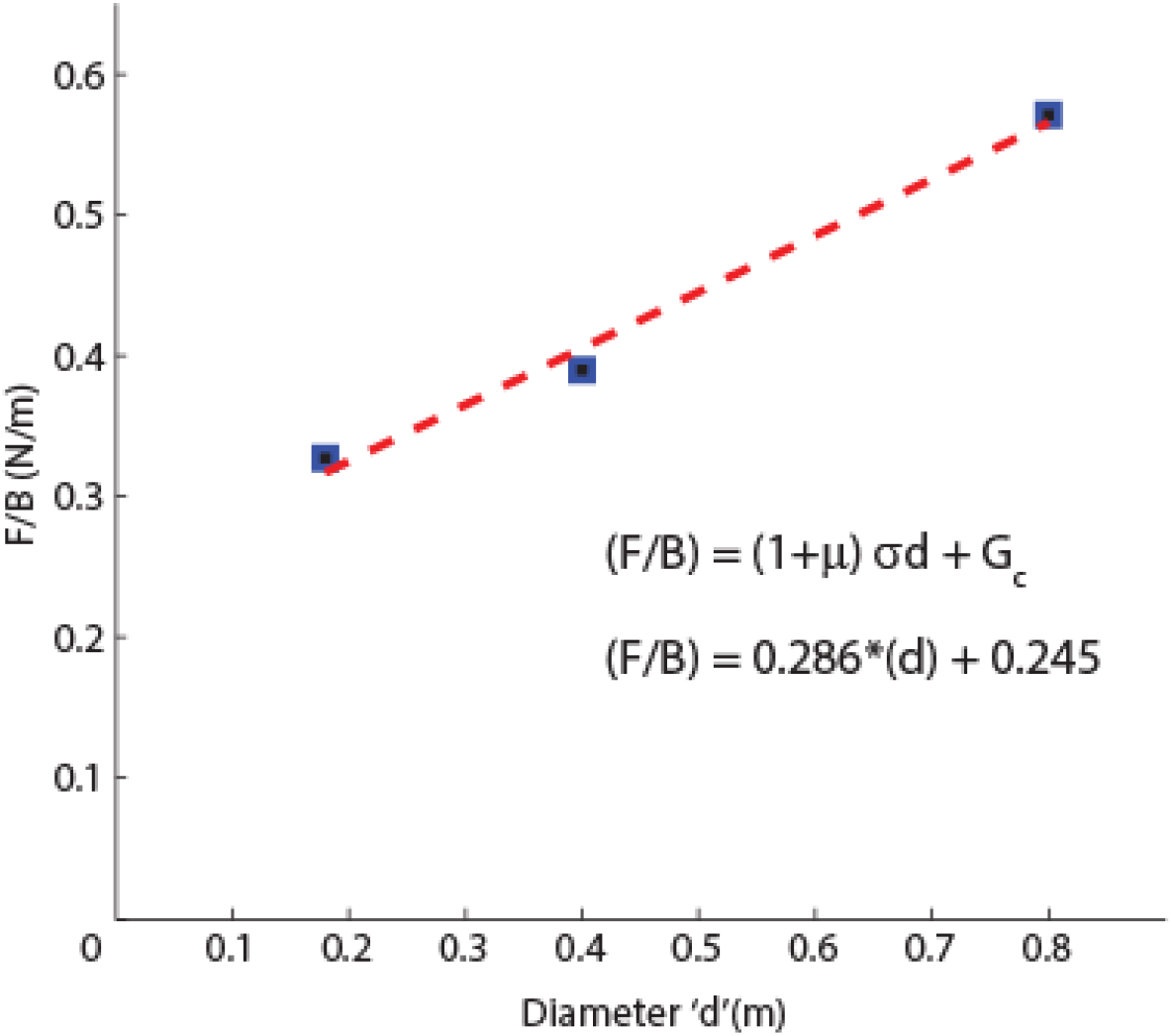
Wire cutting test results from three different diameter wires.

B = sample width, μ = kinematic friction coefficient, *σ* = characteristic stress, and *G_c_* = Griffith critical energy release rate. From the fit, we estimate the values of *G_c_* to be 0.245 J/m^2^. Using this value of *G_c_* and the modulus determined from the compression experiments, we can estimate the stress intensity factor (*K_c_*) using:

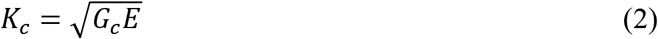

being *E* ≈ 380 kPa the average Young’s modulus of the gelatin. Thus, we get *K_c_* ≈ 0.31 kPa√m.

### 3.2. Modelling

Modelling of the fang-gel interaction was done in three parts. The first part of interaction is the indentation of the gel surface without any insertion (Figure 7, A). The second part of interaction includes sudden piercing of the gel (Figure 7, B), followed by the third interaction that is the continuous insertion of fang into the gel (Figure 7, C). As the fang is pushed more into the gel, there is an increase in the recorded force because of the compression of higher gel volume as a result of the increase in the diameter of the fang from tip to the base. In contrast, the experimental results based on cylindrical needles, the insertion force is almost constant after piercing because of the constant diameter (Mahvash and Dupont, 2010; Okamura et al., 2004). We approximate the fangs as cones and assume the substrate as a linear elastic material to perform modelling of insertion force curve.

**Figure 7.**
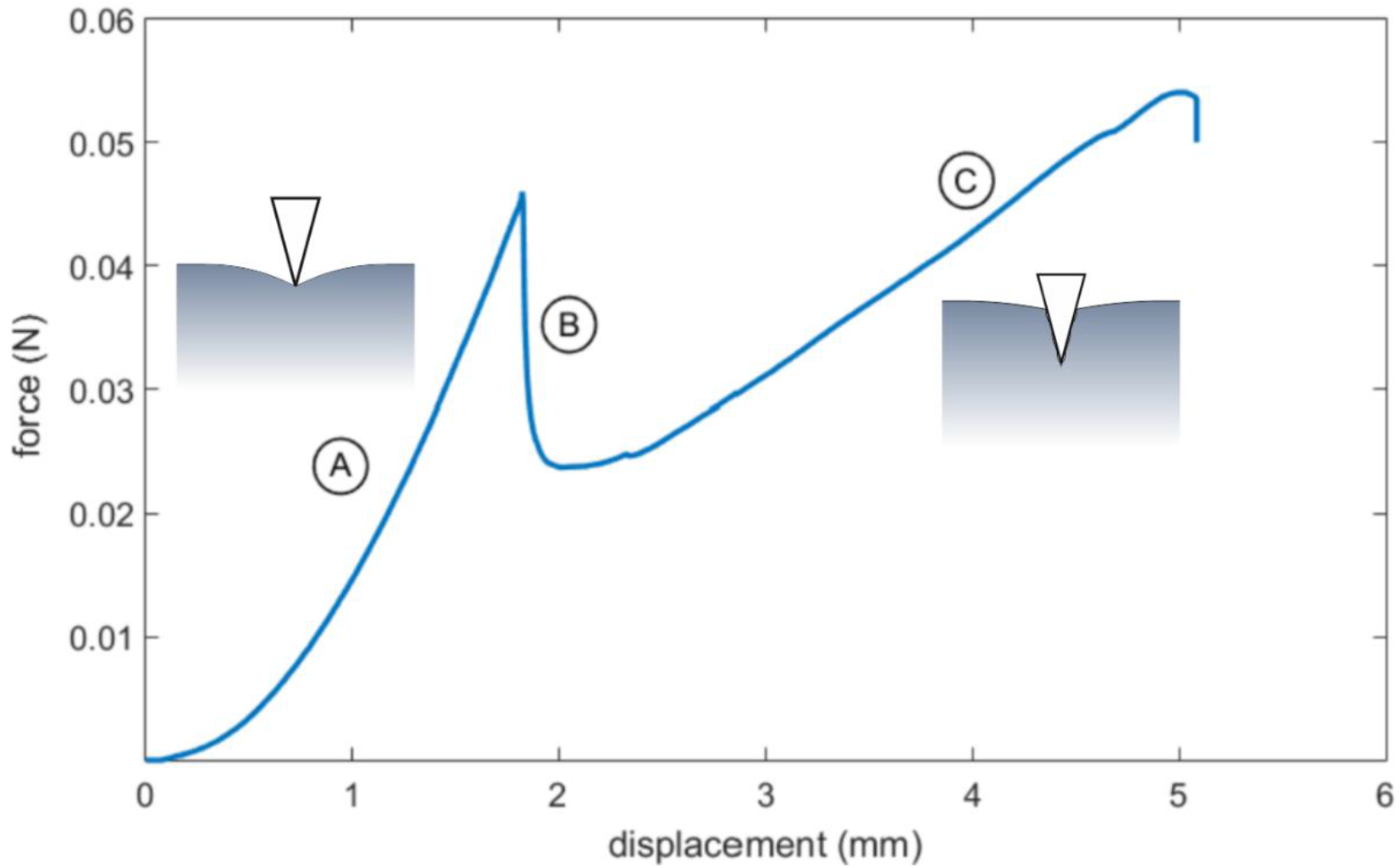
Force-displacement curve with a schematic view of the indentation (left, a) and insertion (right, c) mechanisms. (Example data for the viper fang, *s* = 0.1 mm/s, test no. 1).

#### 3.2.1. Indentation

Let us consider the non-adhesive and frictionless indentation of an elastic half-space by a rigid cone-shaped indenter. The derived relationship between insertion depth (*δ*), contact radius (*a*) and indenter half cone angle (β) can be written as (Sneddon, 1965):

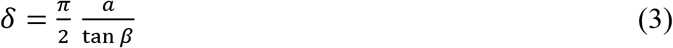

The corresponding indentation force *F_i_* is given by:

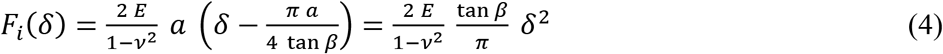

where *E* is the Young’s modulus and *ν* is the Poisson’s ratio of the substrate. We employ equation (4) to fit the experimental data of the indentation part of the curves, by assuming *ν* = 0.5 for the gelatin (Czerner et al., 2015) and *β* = 5.5° for the viper fang and *β* = 8° for the burrowing snake fang, as shown in Figure A1. The deformed shape of the surface outside of the contact area (i.e., for *x* > *a*) is given by (Sneddon, 1965):

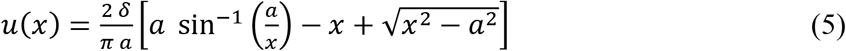

The best fit of the experimental data can be obtained also by adding a linear term (Mahvash and Dupont, 2010; Okamura et al., 2004):

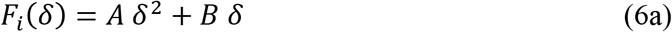

where, *A* and *B* fitting parameters. Although we are not providing an analytical derivation of this additional term, it might be related to one or more of the introduced approximations, i.e., the nonlinear elasticity of the material and the geometry of the fang. The deflection of the surface can also be obtained using a Hertzian fit, i.e.:

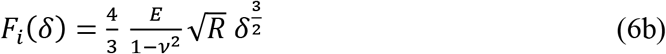

where *R* is the radius of curvature of the tip. Anyway, the assumptions of a Hertz contact are not completely fulfilled here (K.L.Johnson, 1985), in fact the computed Young’s modulus gets underestimated of one order of magnitude and it is not reported here for brevity. The values of Young’s modulus extracted by best-fitting according to Equation (4) are reported in Table 3 and they are in good agreement with the Young’s modulus of the gelatin found through compression testing. The differences observed between the experimental values and those obtained from the model are mostly due the perfect-cone assumption made on the fang shapes. The results do not appear to be strongly affected by the velocity; instead, the effect of the shape of the fangs is more relevant, given the lower values of modulus for the burrowing snake fang.

**Table 3.** Estimated values of various parameters obtained from the fitting of the indentation and insertion parts of the experimental curves (R^2^ values of the fits in brackets).

| sample | Insertion speed ($\nu$)<br>(mm/s) | $E$ (kPa)<br>from Equation<br>(2) | $K_{Ic}$ (kPa $\sqrt{m}$)<br>from Equation<br>(12) | $\sigma$ (kPa)<br>from Equation<br>(12) | $p_{eq}$ (kPa)<br>from<br>Equation (12) |
| --- | --- | --- | --- | --- | --- |
| viper | 0.01 | $244 \pm 36$<br>(0.95) | $1.8 \pm 0.2$<br>(0.99) | $4.9 \pm 1.7$ | $224 \pm 128$ |
| | 0.1 | $229 \pm 36$<br>(0.98) | $1.9 \pm 0.3$<br>(0.99) | $5.1 \pm 2.6$ | $293 \pm 142$ |
| | 1 | $226 \pm 12$<br>(0.98) | $2.2 \pm 0.1$<br>(0.99) | $7.7 \pm 0.3$ | $547 \pm 58$ |
| burrowing snake | 0.01 | $135 \pm 20$<br>(0.98) | $0.9 \pm 0.4$<br>(0.98) | $2.8 \pm 3.5$ | $205 \pm 142$ |
| | 0.1 | $112 \pm 4$<br>(0.99) | $0.7 \pm 0.2$<br>(0.99) | $6.3 \pm 0.9$ | $259 \pm 29$ |
| | 1 | $104 \pm 4$<br>(0.99) | $0.6 \pm 0.2$<br>(0.99) | $6.8 \pm 0.7$ | $555 \pm 52$ |

#### 3.2.2. Insertion

The insertion part of the curves can be described by considering the fracture propagation and the strain energy developed during the progressive piercing. The mechanics of insertion into a soft substrate is driven by the work required to create a unit surface of the crack d*W*_crack_ and the stored strain energy per unit volume d*U*_strain_ (Shergold and Fleck, 2004). Thus, the insertion work expended by the tip must balance the sum of d*W*_crack_ and d*U*_strain_:

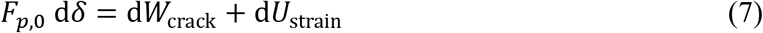

where *F*_*p*,0_ is the insertion force. The infinitesimal lateral area changes of the cone penetrating the substrate of an infinitesimal displacement d*δ* is given by:

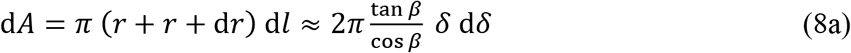

where we have neglected the high-order terms and used *r* = *δ* tan *β* and 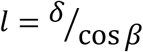. For the volume-related term in Equation (7), i.e. the strain energy, we assume that the stresses arising from the insertion process involve a spherical area around the tip, with radius equal to the insertion depth *δ*. Therefore, we get:

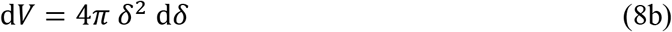

Here we assume that during insertion there is stable crack propagation and the crack maintains a conical shape. Thus, the work required to create an incremental opening of the crack must equal the critical strain energy release rate (Shergold and Fleck, 2004), which, for a mode-I crack opening, is related to the material fracture toughness *K*_I*c*_ through:

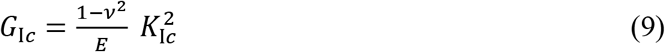

valid for plane-strain conditions. Therefore, by making use of Equation (8a), we get:

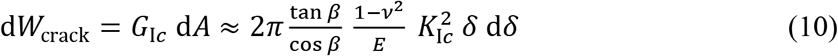

The strain energy, considering the involved volume from Equation (8b), is given by:

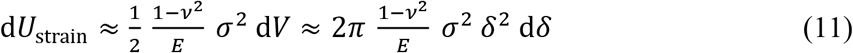

where *σ* can be assumed to be an average stress around the tip during insertion and, again, the plane-strain Young’s modulus is employed.

Finally, we can insert Equations (10) and (11) into Equation (7), obtaining:

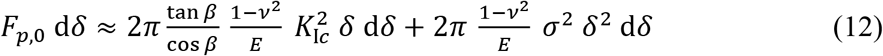

Since Equation (12) must hold for any d*δ*, the force-displacement relationship during insertion is:

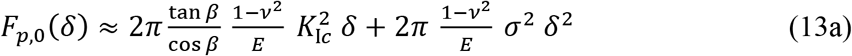

Equation (13a) is obtained by considering the surface of the substrate as flat. Due to friction, this does not happen in experiments, thus the substrate surface remains deflected of a small quantity *α δ_cr_*, with *α* < 1 and *δ_cr_* the critical displacement related to fracture. This effect can be considered, as shown in Figure 6, by:

- considering the effective insertion depth, i.e. applying the change of variable *δ* → *δ* – *αδ_cr_*;
- adding a term due to the indentation of the substrate (up to a depth *α δ_cr_*), as the indentation by an equivalent rigid flat punch of radius *a_cr_*, assumed to be constant in a first approximation. This assumption is reasonable considering that, after the initial quasi-conical shape, the diameter of the fangs becomes almost constant (Fig. A1).

Accordingly, Equation (13a) modifies as follows:

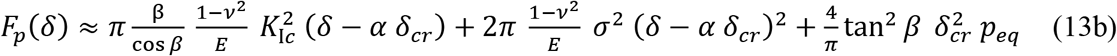

where *p_eq_* is an equivalent pressure related to the flat punch indentation described above, acting on the area 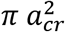, being *a_cr_* related to *δ_cr_* through Equation (2). The result can be considered a measure of the effect of friction and/or adhesion between the fang and the substrate. Rearranging and neglecting high-order terms (i.e. assuming *α*^2^ ≪ 1), we get:

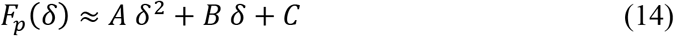

with:

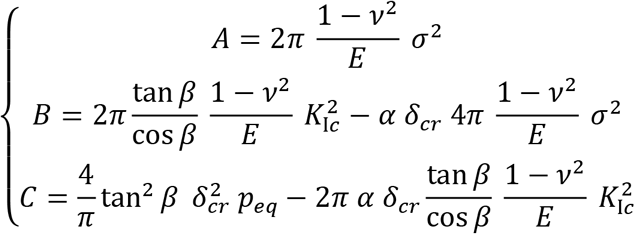

The experimental data was fit to Equation (14) to estimate the fracture toughness of the material *K*_I*c*_, the average stress during insertion *σ* and the equivalent pressure *p_eq_*, as listed in Table 3. The value of the Young’s modulus is taken from the fit of the indentation part of the curves according to Equation (3), while we have chosen *α* ≈ 0.2 from experimental observations. We find that values of *K*_I*c*_ are almost independent from the insertion speed (*v*), while the values of stresses (namely *σ* and *p_eq_*) increased with increase in velocity. Note that the computed fracture toughness of the gelatin as shown in Table 3, is similar to the value obtained through the experimental results of wire cutting tests, especially for the burrowing snake fang. The observed differences can be attributed to inherent assumptions made in modeling and also the differences between the actual shape of the fang and the ideal cone indenter used in modeling.

#### 3.2.3. Fracture

The fracture part of the force-displacement curves is characterised by an instantaneous drop in the force, due to the initial crack formation. We can estimate this force drop by making use of the above-introduced Equations (4) and (14), related to the indentation and insertion part, respectively. In order to employ a formulation of the same type of Equation (14), we introduce the (empirical) dimensionless coefficient *η* ≥ 1, which multiplies the volume-dependent term in the expression of the insertion force (i.e. the term related to strain energy. It is, in other words, a measure to introduce the fracture phenomenon, happening for *δ* = *δ_cr_*. Thus, we get the following semi-analytical expression:

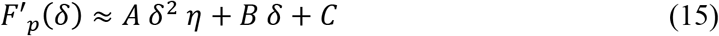

which reduces to Equation (14) for *η* = 1.

Thus, it is possible to extract *η* from Equations (15) and (4) evaluated at *δ* = *δ_cr_* (this latter quantity taken from the experimental data), obtaining:

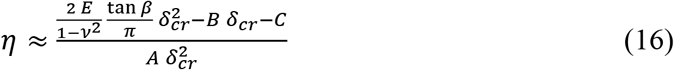

Consequently, the corresponding force drop is found by subtracting Equation (14) from Equation (15) at the critical displacement *δ_cr_*:

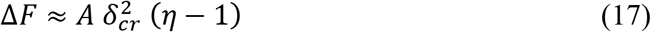

The estimated values of force drop appear to be in the same order of magnitude of those observed in the experiments (Table 4). Differences can be attributed to the introduced assumptions on material and geometry, as already discussed in the derivation of the indentation and insertion laws.

**Table 4.**
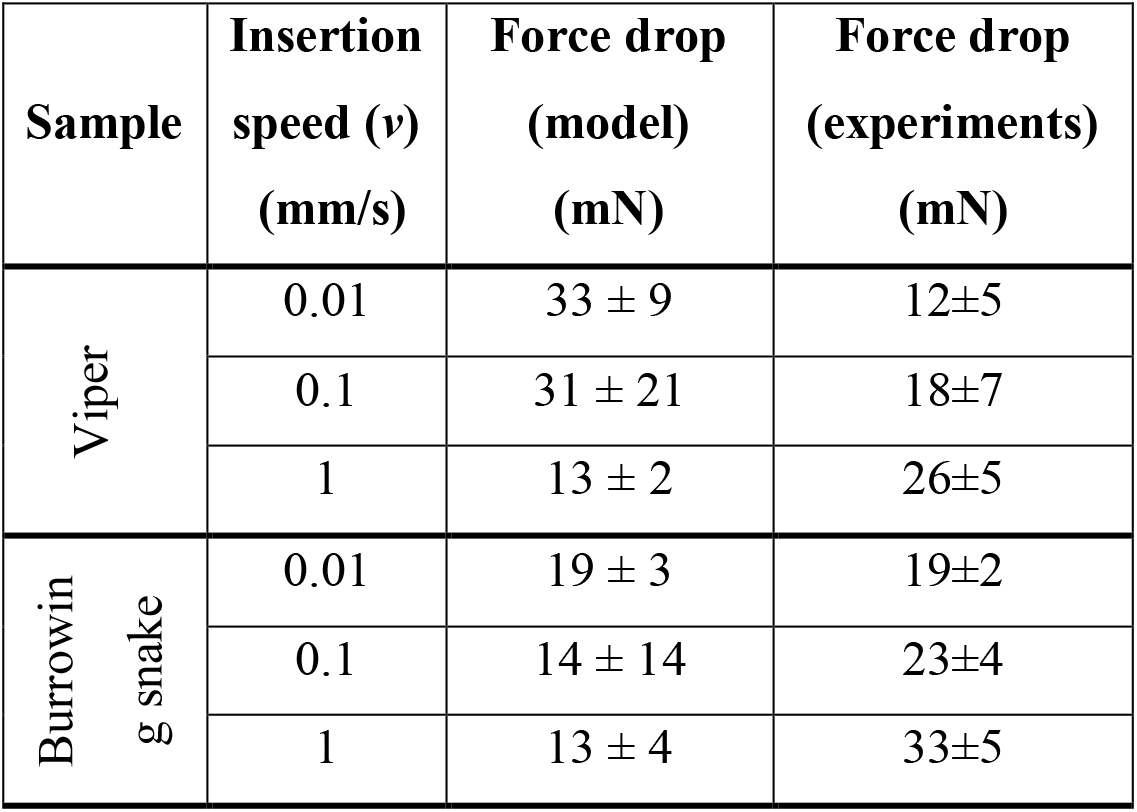
Estimated values of the force drop Δ*F* and comparison with experiments.

| Sample | Insertion speed ($v$)<br>(mm/s) | Force drop (model)<br>(mN) | Force drop (experiments)<br>(mN) |
| --- | --- | --- | --- |
| Viper | 0.01 | $33 \pm 9$ | $12 \pm 5$ |
| | 0.1 | $31 \pm 21$ | $18 \pm 7$ |
| | 1 | $13 \pm 2$ | $26 \pm 5$ |
| Burrowing snake | 0.01 | $19 \pm 3$ | $19 \pm 2$ |
| | 0.1 | $14 \pm 14$ | $23 \pm 4$ |
| | 1 | $13 \pm 4$ | $33 \pm 5$ |

## 4. Conclusions

We compared mechanical properties of fangs of two snake species with different sizes and their also piercing process using gelatine hydrogels. Both the species appear to have similar mechanical properties but there was a difference in the insertion forces owing to the difference in shape of the fang. Our analytical modelling results show that we are able to model the interaction between the fang and the substrate, obtaining a good agreement with the experimental results. Our findings may aid in understanding mechanics and design of bioinspired surgical needles into soft materials such as human skin.

## Acknowledgements

N.M.P. gratefully acknowledges the support of the grants by the European Commission Graphene Flagship Core 2 n. 785219 (WP14 “Composites”) and FET Proactive “Neurofibres” n. 732344 as well as of the grant by MIUR “Departments of Excellence” grant L. 232/2016, ARS01-01384-PROSCAN and PRIN-20177TTP3S. L.K. is supported by Fondazione Caritro under “Laser surface microtexturing for tuning friction”. The authors thank Mirco D’Incau for the help with SEM imaging.

## References

Abràmoff, M. D. and Magalhães, P. J. (2004). Image Processing with ImageJ. Biophotonics Int.

Cundall, D. (2015). Viper fangs: functional limitations of extreme teeth. Physiol. Biochem. Zool. 82, 63–79.

Czerner, M., Fellay, L. S., Suárez, M. P., Frontini, P. M. and Fasce, L. A. (2015). Determination of Elastic Modulus of Gelatin Gels by Indentation Experiments. Procedia Mater. Sci. 8, 287–296.

Jackson, K. (2002). How tubular venom-conducting fangs are formed. J. Morphol. 252, 291–297.

Jackson, K. (2003). The evolution of venom-delivery systems in snakes. Zool. J. Linn. Soc. 137, 337–354.

Jansen van Vuuren, L., Kieser, J. A., Dickenson, M., Gordon, K. C. and Fraser-Miller, S. J. (2016). Chemical and mechanical properties of snake fangs. J. Raman Spectrosc. 47, 787–795.

K.L. Johnson (1985). Contact Mechanics. Cambridge University Press.

Kamyab, I., Chakrabarti, S. and Williams, J. G. (1998). Cutting cheese with wire. J. Mater. Sci. 33, 2763–2770.

Kardong, K. V (2016). “Protovipers” and the Evolution of Snake Fangs Author(s): Kenneth V. Kardong Source: 33, 433–443.

Mahvash, M. and Dupont, P. E. (2010). Mechanics of dynamic needle insertion into a biological material. IEEE Trans. Biomed. Eng. 57, 934–943.

Matushkina, N. and Gorb, S. (2007). Mechanical properties of the endophytic ovipositor in damselflies (Zygoptera, Odonata) and their oviposition substrates. Zoology (Jena). 110, 167–75.

Meyers, M. A., McKittrick, J. and Chen, P.-Y. (2013). Structural biological materials: critical mechanics-materials connections. Science 339, 773–9.

O’Leary, M. D., Simone, C., Washio, T., Yoshinaka, K. and Okamura, a. M. (2003). Robotic needle insertion: effects of friction and needle geometry. 2003 IEEE Int. Conf. Robot. Autom. (Cat. No.03CH37422) 2, 1774–1780.

Okamura, A. M., Simone, C. and O’Leary, M. D. (2004). Force modeling for needle insertion into soft tissue. IEEE Trans. Biomed. Eng. 51, 1707–1716.

Politi, Y., Priewasser, M., Pippel, E., Zaslansky, P., Hartmann, J., Siegel, S., Li, C., Barth, F. G. and Fratzl, P. (2012). A spider’s fang: How to design an injection needle using chitin-based composite material. Adv. Funct. Mater. 22, 2519–2528.

Shergold, O. a. and Fleck, N. a. (2004). Mechanisms of deep insertion of soft solids, with application to the injection and wounding of skin. Proc. R. Soc. A Math. Phys. Eng. Sci. 460, 3037–3058.

Sneddon, I. N. (1965). The relation between load and insertion in the axisymmetric boussinesq problem for a punch of arbitrary profile. Int. J. Eng. Sci. 3, 47–57.

Zahradnicek, O., Horacek, I. and Tucker, A. S. (2008). Viperous fangs: Development and evolution of the venom canal. Mech. Dev. 125, 786–796.

Zhao, Z.-L., Zhao, H.-P., Ma, G.-J., Wu, C.-W., Yang, K. and Feng, X.-Q. (2015). Structures, properties, and functions of the stings of honey bees and paper wasps: a comparative study. Biol. Open 4, 921–928.

